# Kynurenine-AHR signaling sustains a targetable FATP2 dependency in advanced systemic mastocytosis

**DOI:** 10.64898/2026.11.20.766488

**Authors:** Giulia Scorrano, Camille Laisney, Paraskevi Kousteridou, Patricia Kembia Kalombo, Julie Agopian, Rémy Castellano, Armelle Goubard, Arnauld Sergé, Loriane Maillot, Sébastien Létard, Rafaël J Argüello, Yves Colette, Leïla Maouche-Chrétien, Michel Arock, Alberto Orfao, Julien Rossignol, Patrice Dubreuil, Olivier Hermine, Fabienne Brenet

**Affiliations:** Centre de Recherche en Cancérologie de Marseille (CRCM), Inserm U1068, CNRS U7258, Aix-Marseille Univ, Institut Paoli-Calmettes, Marseille, France; Université de Toulouse, Centre de Recherches en Cancérologie de Toulouse (CRCT), Inserm U1037, CNRS U5071, Toulouse, France; Laboratoire d’adhésion et d’inflammation (LAI), Inserm U1067, CNRS U7333, Aix-Marseille Univ UM61, Parc Scientifique et Technologique de Luminy, Marseille, France; Aix-Marseille Univ, CNRS, INSERM, Centre d’Immunologie de Marseille-Luminy (CIML), Marseille, France; Laboratory of Cellular and Molecular Mechanisms of Hematological Disorders and Therapeutic Implications, Inserm U1163, Institut Imagine, Université Paris Cité, Paris, France; Cancer Research Center (IBMCC, USAL-CSIC), Department of Medicine and Cytometry Service (NUCLEUS), CIBERONC, University of Salamanca, Biomedical Research Institute of Salamanca (IBSAL), Salamanca, Spain; Spanish Network on Mastocytosis (REMA), Toledo and Salamanca, Spain; French Reference Center for Mastocytosis (CEREMAST), Groupe Hospitalier Pitié-Salpêtrière, Université Paris Sorbonne, Paris, France

**Keywords:** metabolic adaptation, kynurenine, FATP2 (SLC27A2), KIT inhibition, systemic mastocytosis

## Abstract

Avapritinib and midostaurin have transformed the treatment of advanced systemic mastocytosis (AdvSM), yet residual neoplastic mast cells persist despite sustained KIT inhibition, through mechanisms that remain poorly understood. In a prospective cohort of patients with indolent or advanced SM, plasma metabolomic profiling identified L-kynurenine (L-kyn) as a marker of advanced disease. Transcriptomics analysis of purified bone marrow mast cells, together with single-cell RNA sequencing, further revealed coordinated activation of a tryptophan-kynurenine-aryl-hydrocarbon receptor (AHR) pathway in the neoplastic marrow compartment. In KIT D816V mast cells, L-kyn activated AHR signaling, promoted a metabolic shift toward oxidative phosphorylation and fatty acid oxidation, enhanced AHR-dependent fatty acid uptake, and induced expression of FATP2 (SLC27A2), the most upregulated lipid transporter in AdvSM mast cells. L-kyn also increased plasma membrane-associated cholesterol, partly independently of AHR, whereas AHR knockdown reduced GM1-positive lipid raft abundance. Disruption of membrane sphingolipids and cholesterol impaired KIT and STAT5 phosphorylation, supporting a role for lipid raft integrity in sustaining oncogenic signaling. Neither avapritinib nor midostaurin impaired L-kyn uptake, indicating that this metabolic input persists despite KIT inhibition. Pharmacologic FATP2 inhibition with lipofermata reduced mast cell viability and disease burden in a KIT D816V xenograft model. Pharmacologic or genetic FATP2 blockade synergized with avapritinib and midostaurin in mast cell lines and primary AdvSM cells *ex vivo*, with limited effects on normal CD34+ progenitors., These findings identify FATP2 as a therapeutically actionable metabolic dependency in AdvSM and provide a rationale for combining FATP2 inhibition with KIT-targeted therapy.

## INTRODUCTION

Systemic mastocytosis (SM) is a clonal mast cell (MC) neoplasm ranging from indolent forms with near-normal life expectancy to advanced SM (AdvSM), including aggressive SM (ASM), SM with an associated hematologic neoplasm (SM-AHN), and mast cell leukemia (MCL), which are characterized by organ damage and reduced survival. A gain-of-function KIT mutation, most commonly KIT D816V, drives constitutive kinase signaling in more than 90% of patients and is the principal therapeutic target in SM.^1–6^ Avapritinib, a selective KIT D816V inhibitor, and midostaurin, a multikinase inhibitor, have substantially improved the therapeutic landscape of AdvSM, with overall response rates of approximately 73-75% and 60%, respectively.^3,7–10^ Durable eradication nevertheless remains elusive as complete remission is achieved in only 30 to 36% of avapritinib-treated patients, measurable disease frequently persists during continuous treatment, and both primary and acquired resistance have been reported.^7–9,11–14^ Identifying the mechanisms that enable neoplastic MCs to survive sustained inhibition of oncogenic signaling is therefore a critical unmet need.

The persistence of neoplastic MCs under KIT-directed therapy suggests that cellular fitness is maintained, at least in part, by adaptive programs that kinase inhibition does not adequately suppress, including alternative signaling networks, interactions with the bone marrow microenvironment and metabolic adaptations. In myeloid malignancies, including acute myeloid leukemia (AML), metabolic dependencies have been implicated in therapeutic resistance and have emerged as targets for combination strategies. Whether comparable adaptations support the persistence of neoplastic MCs in AdvSM remains largely unexplored. The kynurenine pathway of tryptophan catabolism generates metabolites that engage the aryl hydrocarbon receptor (AHR),^15,16^ a ligand-activated transcription factor integrating extracellular metabolic cues with cellular metabolic and immune programs across multiple tumor types.^17–20^ Increased circulating kynurenine levels have been reported in mastocytosis and associated primarily with gastrointestinal and neuropsychiatric symptoms.^21,22^ Whether kynurenine contributes directly to neoplastic MC biology has not been investigated. Here, we combined plasma metabolomics, transcriptomic analyses of patient-derived bone marrow MCs, and complementary experimental models to investigate the role of the kynurenine-AHR pathway in AdvSM. We identify FATP2 (SLC27A2) as an AHR-regulated mediator of fatty acid uptake and membrane lipid remodeling that remains accessible to therapeutic targeting during KIT inhibition, and show that pharmacologic or genetic FATP2 blockade enhances the activity of avapritinib and midostaurin in preclinical models.

## METHODS

### Patient samples

Patients with SM were enrolled in the prospective multicenter CEREMAST (French Reference Center for Mastocytosis) study and diagnosed according to WHO criteria, as described.^23^ The plasma metabolomic cohort comprised 13 patients with indolent SM (ISM) and 12 with AdvSM. Bone marrow aspirates for *ex vivo* combination studies were obtained at diagnosis from 4 patients with AdvSM (2 SM-AHN, 2 MCL). All samples were collected after written informed consent in accordance with the Declaration of Helsinki, with approval from the Ethics Committee of Necker Hospital (Paris, France). The CEREMAST/DATAMAST registry was approved by CERAPHP ethics committee (ref 2023-03-01; IORG0010044), complies with CNIL methodology MR-004 and is registered at ClinicalTrials.gov (NCT06186856).

### Cell culture

ROSA cells^24^ were maintained in IMDM with 10% heat-inactivated FCS, penicillin-streptomycin, 2 mM glutamine, and 1 mM sodium pyruvate, with SCF (80 ng/mL) added for ROSA KIT WT. HMC1.1 (KIT V560G) and HMC1.2 (KIT V560G/D816V) cells^25^ were cultured in RPMI 1640 with 10% FCS, antibiotics, and 2 mM glutamine at 3x10^5^ cells/mL and subcultured every 2 to 3 days. Murine bone marrow-derived MCs (BMMCs) were differentiated from KitD814V mice in OptiMEM (10% FCS, antibiotics, 2 mM glutamine, 1 mM sodium pyruvate) with IL-3 (100 ng/mL) for 4 to 6 weeks and used at >90% FcεRI+/KIT+. β-hexosaminidase and cytokine release assays were performed as described.^23^

### siRNA and shRNA transfection

Cells seeded at 0.5x10^5^ cells/mL 24 hours before transfection were electroporated (10^7^ cells, 250 V, 950 µF; Gene Pulser Xcell, Bio-Rad) with 0.4 nmol siControl (#4390844) or siAHR (#4390824, Thermo Fisher), or with shSLC27A2 (TR309351, OriGene), then recovered in complete medium. Knockdown efficiency was confirmed by immunoblot.

### Metabolic fuel dependency (Seahorse and SCENITH)

Fuel dependency was measured on a Seahorse XF analyzer (Agilent) with the Mito Fuel Flex Test kit (UK5099, BPTES, etomoxir). For palmitate oxidation, cells in unbuffered DMEM (2.5 mM glucose, 0.5 mM L-carnitine, pH 7.35) were plated on CellTak-coated plates (45 min, 37°C, CO_2_-free) with etomoxir (4 µM), BSA or BSA-palmitate (200 µM); the oxygen consumption rate (OCR) was recorded at baseline and after oligomycin (1.5 µM), FCCP (0.5 µM), and antimycin A/rotenone (8 µM each). For SCENITH, cells adapted for 1 week to glucose-free DMEM (10 mM galactose, 2 mM glutamine, 1 mM sodium pyruvate) were treated for 48 h with L-kyn (20 µM) with or without CH-223191 (10 µM) and IFNγ (20 ng/mL), and analyzed as described.^26^

### Fatty acid uptake, filipin staining, and L-kyn uptake

For fatty acid uptake, cells stained with live/dead dye (L10120, 15 min) in HBSS (37°C, 5% CO_2_), pre-treated where indicated with lipofermata (10 µM, 1h; MedChemExpress), received BODIPY 500/510 C_1_, C_12_ for 5 min, with oleate as competitor control;^27^ cells were washed, resuspended in PBS/0.5% BSA and analyzed on a BD LSR Fortessa. Filipin staining followed a published protocol.^28^ Cells were fixed (2% PFA, 30 min), quenched (glycine 1.5 mg/mL, 10 min) and stained with filipin (0.3 µg/mL in PBS/10% FBS, 2 h). For live-cell staining, cells adapted for 1h to phenol red-free DMEM F12 with Nutridoma were recorded on an Aurora spectral cytometer (Cytek) for 1min at baseline and for 4 min after addition of filipin (0.3 µg/mL); data were analyzed in FlowJo v10.8.1. L-kyn uptake was measured as described.^29^ Cells in prewarmed HBSS were incubated with L-kyn (200 μM) with or without BCH (10 mM), lysine (5 mM) or leucine for 4 min, fixed (4% PFA, 30 min, dark), washed and acquired on a BD LSR Fortessa (405 nm laser, 450/50 BP filter). For ELISA, cells were treated with L-Trp (75 µM, 30 min); supernatants (1,000g, 20 min) and lysates (freeze-thaw, 5,000g, 5 min) were assayed with the Human Kynurenine ELISA Kit (KTE61870, Abbkine).

### RT-qPCR and immunoblotting

RNA was extracted with the RNeasy Mini kit (Qiagen) and cDNA synthesized from 200 ng RNA (AffinityScript Multiple Temperature Kit, Agilent). qPCR used SsoAdvanced Universal SYBR Green Supermix (Bio-Rad) with 0.5 μM primers: *SLC27A2* (Fwd GGTGATATTTTCCACTCAG, Rev TCATAGATGCATATGTCCCC); *CD36* (Fwd AGCTTTCCAATGATTAGACG, Rev GTTTCTACAAGCTCTGGTC); *HPRT1* (Fwd ATAAGCCAGACTTTTTGG, Rev ATAGGACTCC AGATTTTCC). For immunoblotting, cells were lysed in RIPA buffer and 30 µg of protein was resolved on 10% SDS-PAGE. Membranes blocked in 5% BSA/TBST were incubated overnight at 4°C with primary antibodies, then with HRP-conjugated secondary antibodies for 1h, and developed by chemiluminescence (ImageQuant 800). For nuclear and cytoplasmic fractionation, 10^7^ ROSA cells treated with L-kyn (20 μM, 30 min) were lysed with NE-PER reagents (78833, Thermo Fisher). Antibodies: FATP2 (1:500, Santa Cruz); phosphoKIT Y568-570 (1:500, Invitrogen); KIT (1:1000, Cell Signaling); phosphoSTAT5 Y694, STAT5, phosphoSTAT3, AHR, and ERK2 (1:1000, Cell Signaling); STAT3, and CD45 (1:1000, Santa Cruz); α-Tubulin (1:1000, Invitrogen), and lamin B1 (1:1000, Invitrogen).

### Cell viability and drug synergy

Cells pre-treated overnight with L-kyn (20 µM) received lipofermata (0.6 µM, 4h) followed by KIT inhibitors (250 nM, 72h). Viability was measured by AlamarBlue (Sigma-Aldrich; CLARIOstar Plus). Synergy was assessed from dose-response matrices in SynergyFinder 2.0 using the ZIP model (scores below -10 indicate antagonism, -10 to 10 additivity, and above 10 synergy).^30^

### Membrane lipid analyses

For GM1 quantification, fixed (2% PFA, 30 min) siCTRL and siAHR HMC1.2 cells were stained with Alexa Fluor 594-conjugated cholera toxin subunit B (CT-B; 30 min, 4°C) and mounted on poly-L-lysine IBIDI microslides. Mid-cell epifluorescence images were smoothed (15-pixel median filter, 0.11 µm/pixel), segmented by moment-based thresholding, and refined by morphological dilation and erosion (80 pixels) using a custom MATLAB script (courtesy of D. Martín)^31^; objects smaller than 10^4^ pixels were excluded. Membrane-ring fluorescence was quantified after subtraction of an inner mask (eroded by 80 pixels) and of background. For raft destabilization, HMC1.2 cells were treated with myriocin (25 µM, 48h), transferred to DMEM-F12 with Nutridoma, then co-treated with myriocin (25 µM) and zaragozic acid (10 µM) for 16h; where indicated, SCF was added for 5 min before lysis.

### Transcriptomic and single-cell analyses

Differential expression between AdvSM vs ISM bone marrow MCs was assessed with limma^32^ in a published dataset of sorted MCs.^40^ Heatmaps used pheatmap v1.0.12 and RColorBrewer v1.1.3; gene-set overlaps (curated metabolic genes, AHR targets,^33^ and KEGG lipid metabolism) used venn v1.11; GSEA used fgsea (1,000 permutations, MsigDB/KEGG signatures); volcano and PCA plots used ggplot2 v3.4.3 and ggbiplot v0.55. Public single-cell RNA-sequencing data (GEO GSE249445^34^; GSM7947695–GSM7947697, GSM7947680, GSM7947683, GSM7947686) were analyzed in Seurat v5.3.1^35^. Cells were quality-filtered, SCTransform-normalized, integrated across samples^36^, and visualized by UMAP; clusters were defined with the Louvain algorithm, and signature scores computed with AddModuleScore^37^ were compared between SM-AHN and healthy donor samples by Wilcoxon test.

### Xenogeneic transplantation model

ROSA KIT D816V-GFP cells transduced with a lentiviral Gaussia luciferase (Gluc)/CFP construct^38^ were injected into the tail vein of sublethally irradiated (1.5Gy) NOD-SCID IL-2Rγ^-/-^ (NSG) mice 24h after irradiation. Three weeks later, mice were randomized to vehicle or lipofermata (2 mg/kg intraperitoneally, 5 days per week, n=10 per group). Engraftment was monitored by plasma Gluc activity and flow cytometry of GFP+ cells.^38,39^ Studies complied with French regulations under APAFIS authorization N°379212022070413517007.

### Statistical analysis

Statistical analyses used GraphPad Prism 8.0.1 and R. Data are shown as mean ± SEM or SD, as indicated. Two groups were compared by unpaired two-tailed Student’s t tests or the corresponding nonparametric tests, and multiple groups by one-way ANOVA with Tukey’s post hoc correction.

## RESULTS

### Circulating L-kynurenine marks advanced disease and reflect activation of a tryptophan-AHR metabolic program in the neoplastic bone marrow

We performed an extended analysis of our previously reported plasma metabolomic profiles from a prospective CEREMAST cohort (13 ISM and 12 AdvSM) in which L-kyn had emerged as a candidate discriminator alongside N-acetylglucosamine (GlcNAc).^23^ L-kyn is a major intermediate of the kynurenine pathway, the predominant route of tryptophan catabolism and a source of NAD+ precursors (Figure S1A). Among the ten most dysregulated metabolites, two inversely correlated clusters emerged. L-kyn, quinolinate, cholesterol, pyridoxine, and GlcNAc were increased in AdvSM, whereas guanosine and cytidine monophosphates, dephospho-CoA, dihydroxyacetone phosphate, and glyceraldehyde 3-phosphate were decreased (Figure 1A). L-kyn levels were significantly higher in AdvSM than in ISM and clustered with quinolinate and pyridoxine, a cofactor for several enzymes of the kynurenine pathway, consistent with enhanced pathway activity in advanced disease (Figure 1B). L-kyn discriminated advanced from indolent disease with an area under the curve (AUC) of 0.94, outperforming quinolinate (Figure 1C, S1B-D).

**Figure 1:**
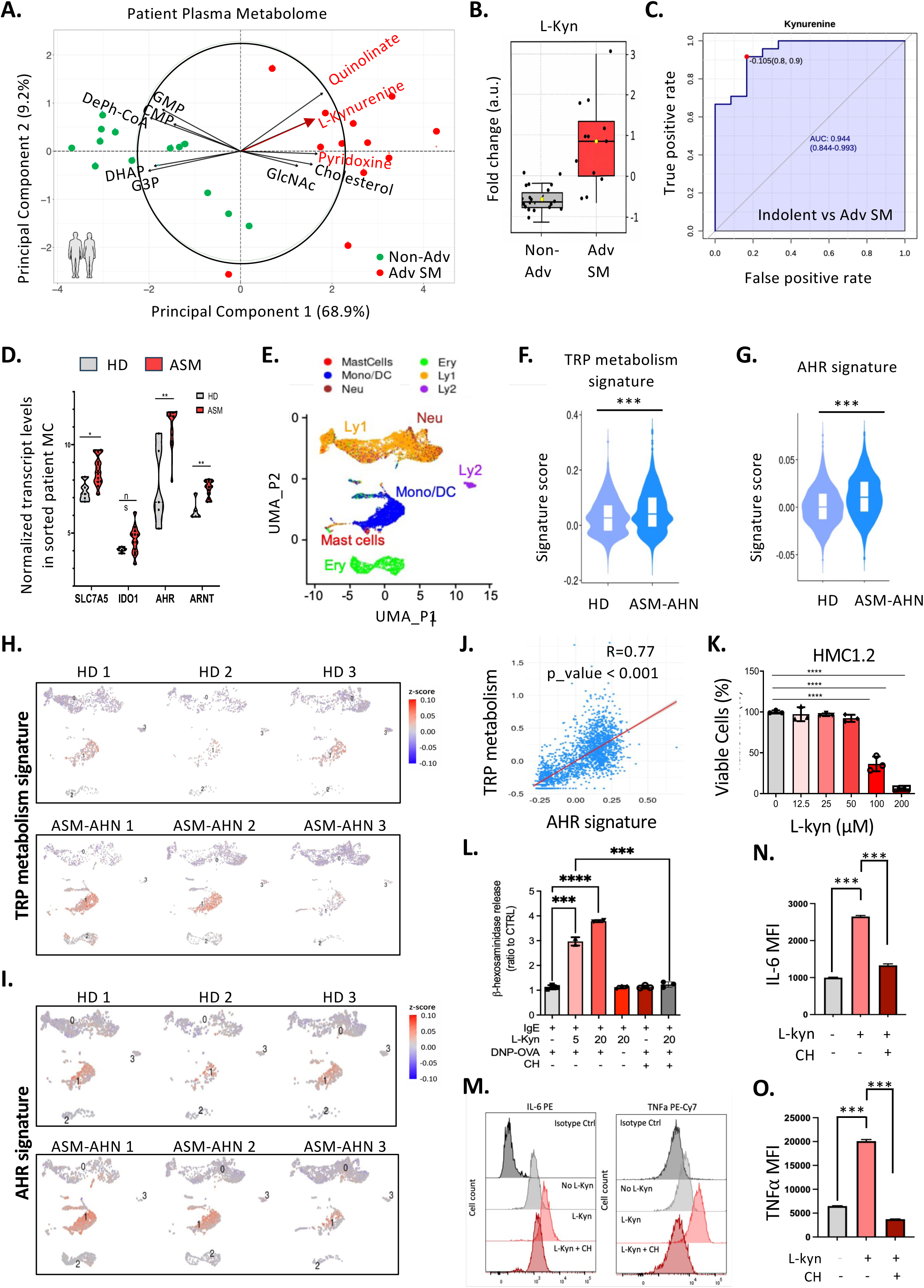
Plasma kynurenine distinguishes advanced systemic mastocytosis and modulates mast cell activation. (A) Correlation plot of the most dysregulated plasma metabolites in indolent (ISM) and advanced systemic mastocytosis (AdvSM), highlighting the clustering of L-kyn with quinolinate, pyridoxine, cholesterol, and N-acetyl-D-glucosamine (B) Relative plasma L-kyn levels in patients with ISM (n=13) and advanced SM (n=12). (C) Receiver operating characteristic (ROC) curve assessing the ability of plasma L-kyn to discriminate AdvSM from ISM. (D) Expression of genes involved in the tryptophan-kynurenine-AHR pathway genes in sorted bone marrow MCs from healthy donors and patients with SM. (E-J) Single-cell RNA-seq analyses showing enrichment of tryptophan metabolism and AHR transcriptional signatures in advanced disease and their correlation within a bone marrow niche enriched for monocyte and MCs. (K) Viability of HMC1.2 cells following exposure to increasing concentrations of L-kyn. (n=3). (L) β-hexosaminidase release by *Kit*D814V bone marrow-derived mast cells (BMMCs) stimulated with IgE/antigen in the presence or absence of L-kyn and the AHR inhibitor CH-223191. (n=3) (M-O) Intracellular IL-6 and TNF-α levels in stimulated BMMCs treated with vehicle, L-kyn or CH-223191 (n=3). Data are presented as mean ± SEM from at least three independent experiments, where applicable. *P < 0.05; **P < 0.01; ***P < 0.001; ****P < 0.0001; ns, not significant.

Because circulating metabolites may originate from any tissue, we asked whether the pathway is engaged within the neoplastic compartment itself. Transcriptomic profiling of sorted human bone marrow MCs^40^ revealed increased expression of the tryptophan/L-kyn transporter LAT1, the rate-limiting enzyme IDO1, AHR, and its partner ARNT in SM relative to healthy donors (Figure 1D). We then analyzed single-cell RNA sequencing of bone marrow samples from patients and healthy donors, annotated using established lineage markers.^34,41^ This identified a monocyte/MC-enriched population with coordinated expression of tryptophan-kynurenine and AHR signatures (Figure 1E-G, S2A), both significantly enriched in SM-AHN relative to healthy donors (Figure 1H-J). Because IDO1 is also expressed by monocytes, these findings establish activation of the tryptophan-kynurenine-AHR program within the neoplastic bone marrow compartment without excluding a contribution from non-MC populations.

We next examined whether L-kyn directly modulates MC function. At 20 µM, the lowest concentration sufficient to induce AHR nuclear translocation (Figure S2C-G), L-kyn did not affect the viability of murine or human MCs (Figure 1K, S2B). However, it potentiated β- hexosaminidase release and IL-6 and TNF-α production after IgE/antigen stimulation of kitD814V BMMCs, an effect abrogated by the AHR antagonist CH-223191 (Figure 1 L-O). L-kyn alone did not trigger degranulation, indicating that it primes rather than activates MCs and providing a potential link with the mediator-related symptoms of SM.

### Kynurenine-AHR signaling shifts neoplastic mast cell metabolism toward mitochondrial oxidative metabolism and fatty acid oxidation

Given the role of the kynurenine pathway in NAD+ precursor production, we investigated its contribution to the bioenergetic phenotype of neoplastic MCs. Gene set enrichment analysis of bone marrow MC transcriptomes showed enrichment of oxidative phosphorylation (OXPHOS) signatures in AdvSM compared with ISM, and single-cell analysis identified an OXPHOS-high population associated with increased activity of the tryptophan metabolic program (Figure 2A-D).^40^

**Figure 2:**
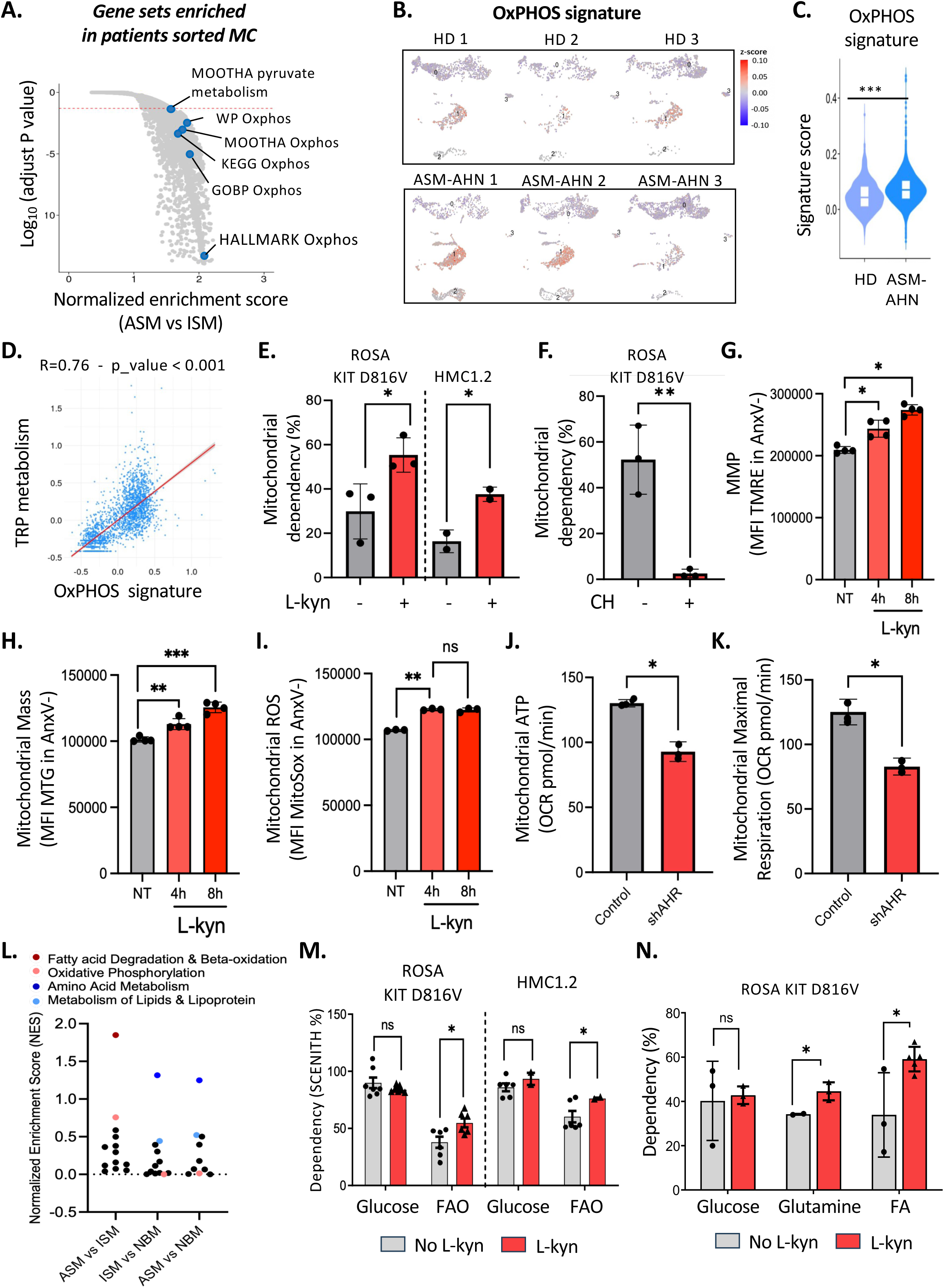
Kynurenine-AHR signaling promotes oxidative metabolism and fatty acid oxidation in neoplastic mast cells. (A) Gene set enrichment analysis (GSEA) showing enrichment of oxidative phosphorylation (OXPHOS)-related gene signatures in MCs from patients with AdvSM compared with ISM. (B-D) Single-cell RNA sequencing analyses demonstrating increased OXPHOS program activity in advanced disease and its correlation with the tryptophan metabolism signature. (E-F) SCENITH-based assessment of mitochondrial dependency in neoplastic MCs following L-kyn exposure, with or without AHR inhibition (n=3). (G-I) Assessment of mitochondrial membrane potential (n=4), mitochondrial mass (n=4), and mitochondrial reactive oxygen species (ROS) in viable neoplastic MCs following L-kyn treatment (n=3). (J-K) Mitochondrial ATP content and maximal oxygen consumption rate (OCR) in control and AHR-deficient cells (n=3), demonstrating partial reversal of the L-kyn-induced bioenergetic phenotype. (L) mitoXplorer-based analysis of mitochondrial transcriptional programs in patient-derived MCs, highlighting preferential enrichment of fatty acid degradation and β-oxidation pathways in AdvSM. (M) SCENITH-based assessment of metabolic dependency demonstrating increased reliance on fatty acid oxidation (FAO) following L-kyn exposure (n=7). (N) Seahorse-based palmitate oxidation assay demonstrating an increased contribution of long-chain fatty acids to basal mitochondrial respiration in KIT D816V neoplastic MCs (n=3). Data are represented as mean ± SEM from at least three independent experiments, where applicable. *P < 0.05; **P < 0.01; ***P < 0.001; ns, not significant.

We then assessed metabolic dependencies using SCENITH.^26^ L-kyn exposure increased mitochondrial dependency in both ROSA KIT D816V and HMC1.2 cells, an effect abolished by CH-223191 (Figure 2E-F, S4A-C). This was accompanied by increased mitochondrial membrane potential, mass, and reactive oxygen species (ROS) in viable cells (Figures 2G-I). L-kyn also increases intracellular ATP content and the maximal oxygen consumption rate (OCR), effects only partially reversed by AHR knockdown (Figures 2J, K) identifying AHR as a major but not exclusive determinant of the L-kyn-induced bioenergetic response.

To define the substrates supporting this mitochondrial activity, analysis of patient-derived MC transcriptomes using MitoXplorer^44^ revealed progressive enrichment of fatty acid degradation and β-oxidation genes from healthy donors to ISM and AdvSM (Figure 2L). Accordingly, SCENITH showed increased dependence on mitochondrial fatty acid oxidation after L-kyn exposure in both models (Figure 2M). Seahorse analysis identified palmitate as the predominant contributor to basal respiration in L-kyn-treated ROSA KIT D816V cells (Figure 2N). Kynurenine-AHR signaling therefore promotes mitochondrial oxidative metabolism with a pronounced reliance on fatty acids, suggesting an increased requirement for exogenous lipid acquisition.

### FATP2 emerges as an AHR-associated lipid transporter upregulated in advanced disease and mediates kynurenine-induced fatty acid uptake

We intersected a curated list of metabolic enzymes and transporters with a published AHR target gene set.^33^ Restricting the analysis to transcripts differentially expressed between AdvSM and ISM MCs yielded 365 candidate genes (Figure 3A). Functional enrichment highlighted long-chain fatty acid metabolism and β-oxidation, sphingolipid, and phosphatidylinositol metabolism, with coordinated downregulation of ion transport and glucose metabolism (Figure S3A). RNA sequencing of neoplastic MCs exposed to L-kyn reproduced this signature (Figure S3B-C).

**Figure 3:**
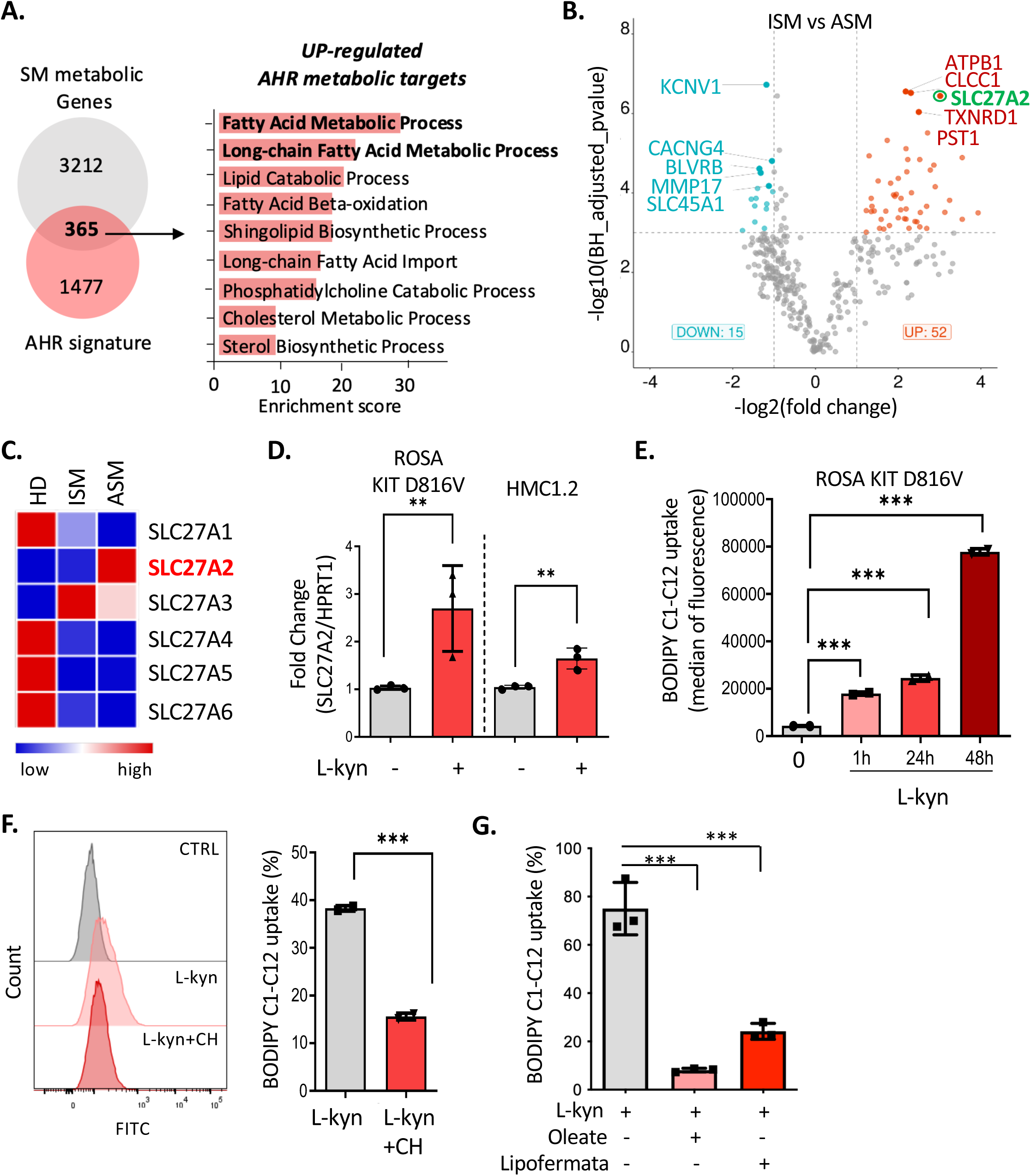
FATP2 is an AHR-associated fatty acid transporter upregulated in advanced systemic mastocytosis. (A) Transcriptomic analysis identifying an AHR-associated metabolic program enriched in SM-associated metabolic genes. (B) Differential gene expression analysis identifying SLC27A2/FATP2 among AHR-associated genes enriched in MCs from patients with AdvSM compared with ISM. (C) Heatmap showing expression of FATP family members across healthy donor, ISM, and AdvSM MC samples, highlighting preferential FATP2 expression in advanced disease. (D) RT-qPCR analysis of SLC27A2 expression following L-kyn exposure in HMC1.2 and ROSA KIT D816V cells. (n=3) (E) Time-course analysis of fluorescent fatty acid analog BODIPY C1-C12 uptake in neoplastic MCs following L-kyn treatment. (n=3) (F) Effect of AHR inhibition with CH-223191 on L-kyn-induced fatty acid uptake in HMC1.2 cells. (n=3). (G) Effect of pharmacological FATP2 inhibition with lipofermata on BODIPY-labeled fatty acid uptake in HMC1.2 cells. (n=3). Data are represented as mean ± SEM from at least three independent experiments, where applicable. *P < 0.05; **P < 0.01; ***P < 0.001; ns, not significant.

Among fatty acid transport proteins, SLC27A2, encoding fatty acid transport protein 2 (FATP2), was the most strongly upregulated and the most highly expressed SLC27A family member in AdvSM relative to ISM and healthy donors (Figure 3B-C). FATP2 is well suited to couple lipid acquisition with intracellular utilization, acting both as a fatty acid transporter and as a very-long-chain acyl-CoA synthetase that activates imported fatty acids, frequently in cooperation with CD36.^45,46^

L-kyn increased SLC27A2 transcript levels in HMC1.2 and ROSA KIT D816V cells (Figure 3D), together with a time-dependent increase in uptake of the fluorescent fatty acid analog BODIPY C1-C12 (Figure 3E). AHR inhibition with CH-223191 markedly attenuated this increase, (Figure 3F), and FATP2 inhibition with lipofermata reduced fatty acid uptake in HMC1.2 cells (Figure 3G), identifying FATP2 as a critical component of the lipid acquisition machinery engaged by kynurenine signaling.

### Kynurenine-driven lipid remodeling promotes membrane cholesterol enrichment and supports KIT-associated lipid raft integrity

Imported fatty acids also contribute to membrane lipid synthesis. Given the enrichment of sphingolipid, glycerophospholipid, ether lipid, and cholesterol pathways within the AHR-associated program (Figure 4A) and the increased circulating cholesterol in AdvSM (Figure 1A), we asked whether kynurenine signaling alters membrane lipid composition.

**Figure 4:**
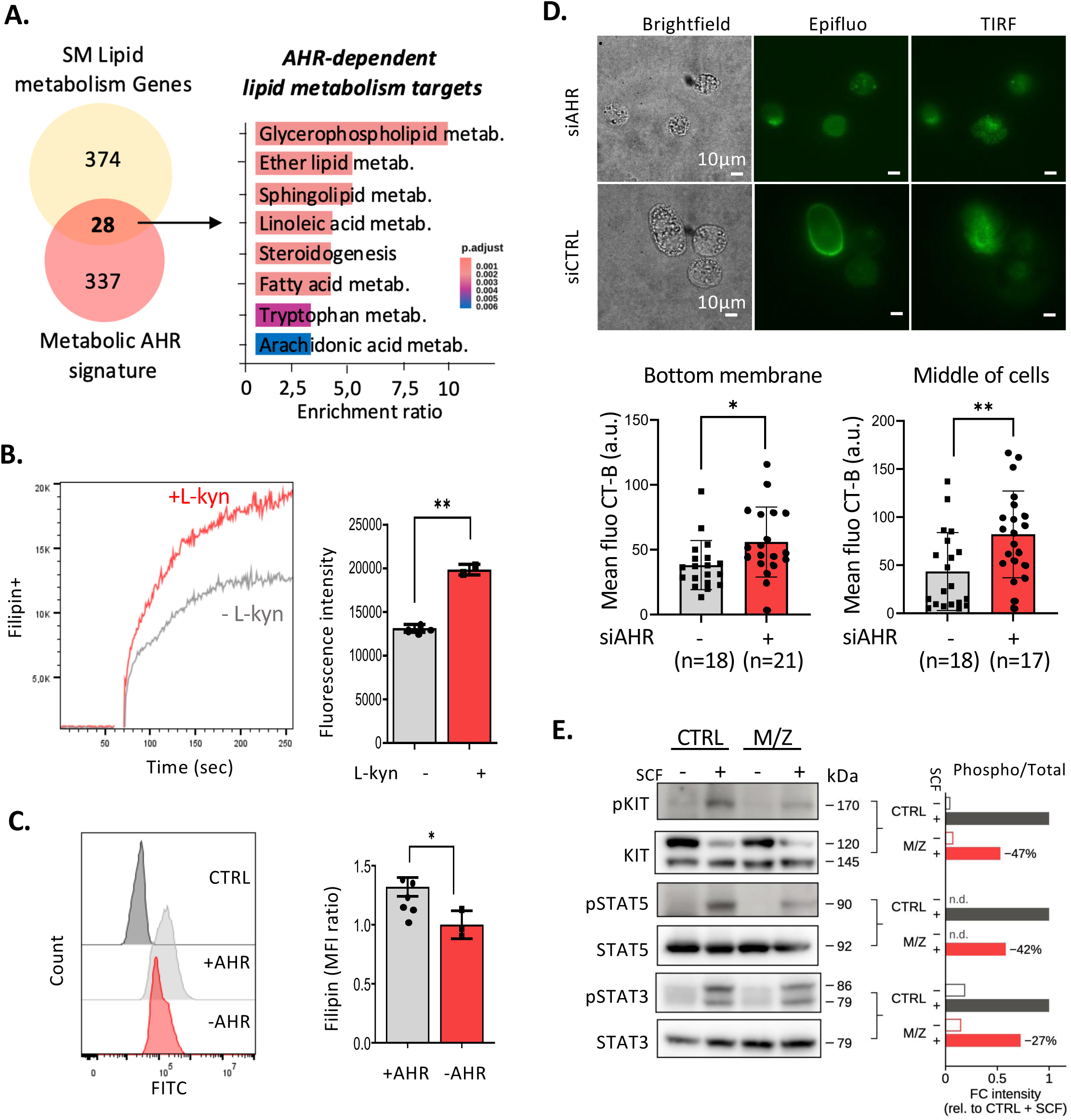
Kynurenine-AHR signaling remodels membrane lipid composition to sustain oncogenic KIT signaling. (A) Transcriptomic analysis identifying an AHR-associated gene program enriched in lipid metabolism pathways (B) Live-cell filipin staining kinetics demonstrating increased plasma membrane cholesterol accumulation following L-kyn treatment (n=3). (C) Filipin staining and quantification in control and AHR-deficient neoplastic MCs, demonstrating partial AHR dependence of cholesterol accumulation (n=3). (D) Cholera toxin B staining and quantification of GM1-positive membrane domains, demonstrating reduced lipid raft content following AHR knockdown. (E) Immunoblot analysis of KIT pathway signaling following lipid raft disruption with myriocin and zaragozic acid, showing reduced KIT and downstream STAT5 phosphorylation (n=2); (pKIT Y568/Y570, pSTAT5 Y694, pSTAT3 Y705). Data are represented as mean ± SEM from at least three independent experiments, where applicable. *P < 0.05; **P < 0.01; ***P < 0.001; ns, not significant.

Filipin staining revealed an increase in free cholesterol accumulation at the plasma membrane following L-kyn exposure, an effect that was partially dependent on AHR (Figure 4B-C). Part of this response persisted after AHR knockdown, suggesting additional transcription-independent contributions by kynurenine pathway metabolites. AHR knockdown further reduced GM1-positive microdomains, at the plasma membrane and in intracellular compartments, assessed by cholera toxin B (CT-B) fluorescence (Figure 4D).

Because KIT signaling is organized within lipid raft microdomains, whose lipid composition is essential for the assembly and function of membrane-associated signaling complexes,^47,48^ we assessed the contribution of membrane lipids to KIT activation. Combined inhibition of sphingolipid and cholesterol biosynthesis with myriocin and zaragozic acid markedly reduced SCF-induced phosphorylation of KIT Y568/Y570 and STAT5 Y694, with a more modest effect on STAT3 phosphorylation at Y705 (Figure 4E). Membrane lipid composition is therefore required for efficient KIT signaling, supporting a model in which kynurenine-associated lipid remodeling maintains signaling-competent raft domains.

### Pharmacological FATP2 inhibition preferentially targets neoplastic mast cells and reduces disease burden *in vivo*

We next evaluated the therapeutic tractability of FATP2 by comparing two structurally distinct inhibitors. Lipofermata^49,50^ induced concentration-dependent cell death with an IC50 of 1.6 μM, whereas grassofermata^51^ was less potent (IC50, 4.6 μM). Under identical experimental conditions, the IC50 values for avapritinib and midostaurin were 0.9 μM and 1 μM, respectively (Figure 5A). Lipofermata was active across all human neoplastic MC lines irrespective of KIT mutational status, whereas HS5 stromal cells, A-498 renal carcinoma, and BT-20 breast cancer cells were substantially less sensitive (Figure 5B), and it was the most potent of the metabolic inhibitors tested in parallel (Figure S5). Activity across distinct KIT genotypes suggests that fatty acid uptake is a broad dependency of neoplastic MCs rather than a vulnerability restricted to KIT D816V-driven disease.

**Figure 5:**
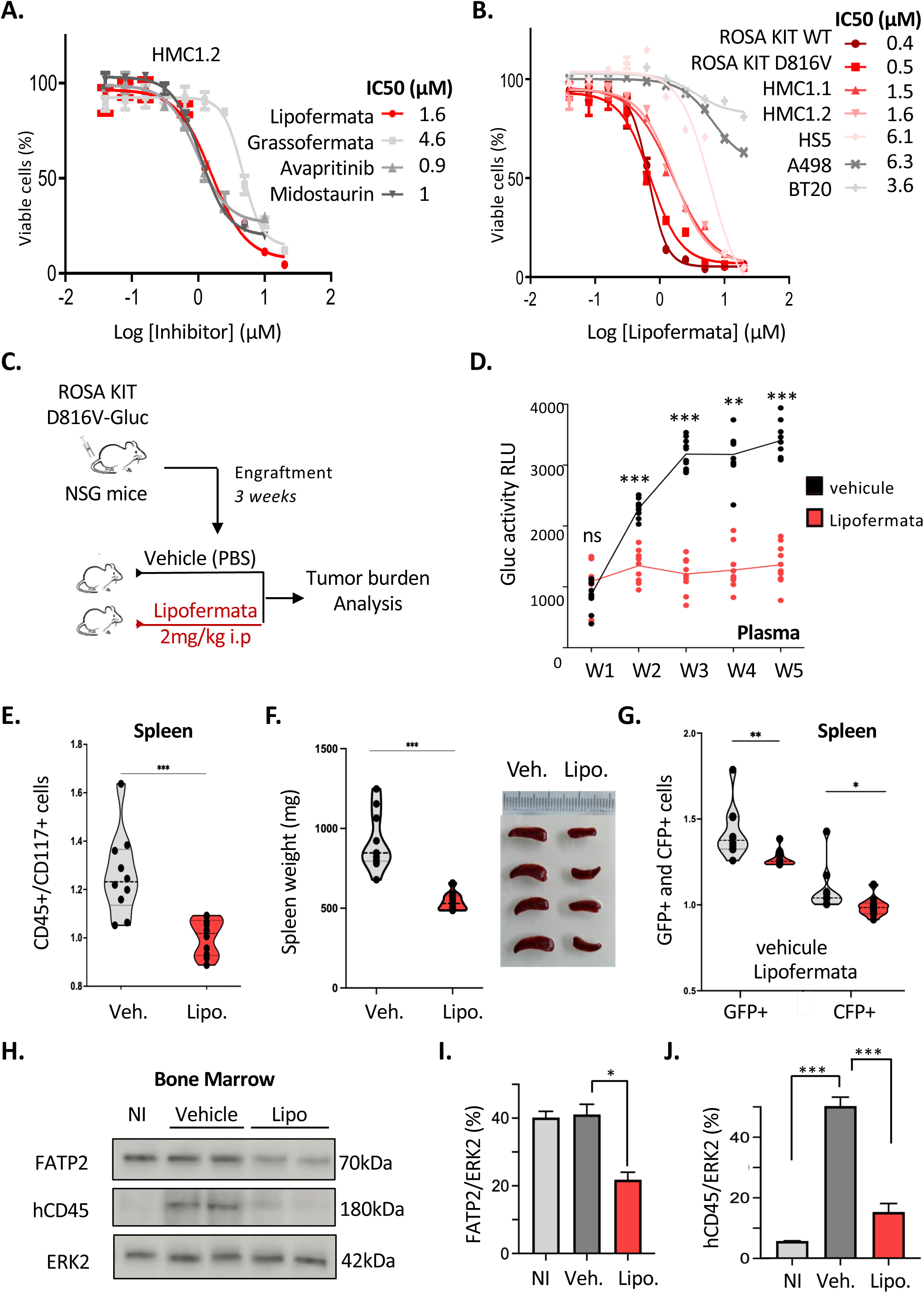
FATP2 inhibition reduces disease burden in KIT D816V xenograft model. (A) Dose-response curves comparing the effects of FATP2 and KIT inhibitors on HMC1.2 cell viability. (B) Comparative sensitivity of neoplastic MC lines and non-MC cell lines to FATP2 inhibition with lipofermata. (C) Schematic representation of the experimental design and treatment schedule for lipofermata administration in the KIT D816V xenograft model (n=10 mice per group). (D) Longitudinal assessment of disease burden by plasma Gaussia luciferase activity in vehicle- and lipofermata-treated mice. (E-G) Quantification of engrafted neoplastic cells in the spleen based on CD45, CD117, GFP, and CFP expression. (H-J) Assessment of bone marrow disease burden following lipofermata treatment, including FATP2 and human CD45 expression by immunoblotting. Each replicate corresponds to one pool of five mice (NI, non-injection; veh, vehicule; lipo, lipofermata). For in vivo experiments (C-J), each symbol represents an individual mouse (n=10 per group). Data are represented as mean ± SEM from at least three independent experiments, where applicable. *P < 0.05; **P < 0.01; ***P < 0.001; ns, not significant.

We then tested this dependency *in vivo* in a xenograft model of aggressive SM/MCL-like disease, in which ROSA KIT D816V cells expressing Gaussia luciferase (Gluc) engraft in sublethally irradiated NSG mice, allowing longitudinal assessment of disease burden by plasma Gluc activity and GFP/CFP labeling.^24,38^ Treatment with lipofermata (2 mg/kg intraperitoneally, 5 days per week) was initiated 3 weeks after engraftment (Figure 5C). Lipofermata-treated animals exhibited significantly lower plasma Gluc activity from the second week of treatment onward (Figure 5D). This was accompanied by decreased splenic CD45+CD117+, GFP+, and CFP+ cells (Figure 5E-G) and by reduced FATP2 and human CD45 signals in the bone marrow (Figure 5H-J), establishing FATP2-dependent lipid import as a therapeutically exploitable vulnerability in advanced SM.

### The kynurenine-FATP2 axis persists despite KIT inhibition, and FATP2 blockade enhances sensitivity to avapritinib and midostaurin

Because residual disease persists under KIT-targeted therapy, we asked whether the kynurenine-FATP2 axis remained functional during KIT inhibition. Neither avapritinib nor midostaurin reduced L-kyn uptake by neoplastic MCs (Figure 6A), indicating that kynurenine acquisition is maintained independently of KIT kinase activity and remains accessible despite suppression of KIT signaling.

**Figure 6:**
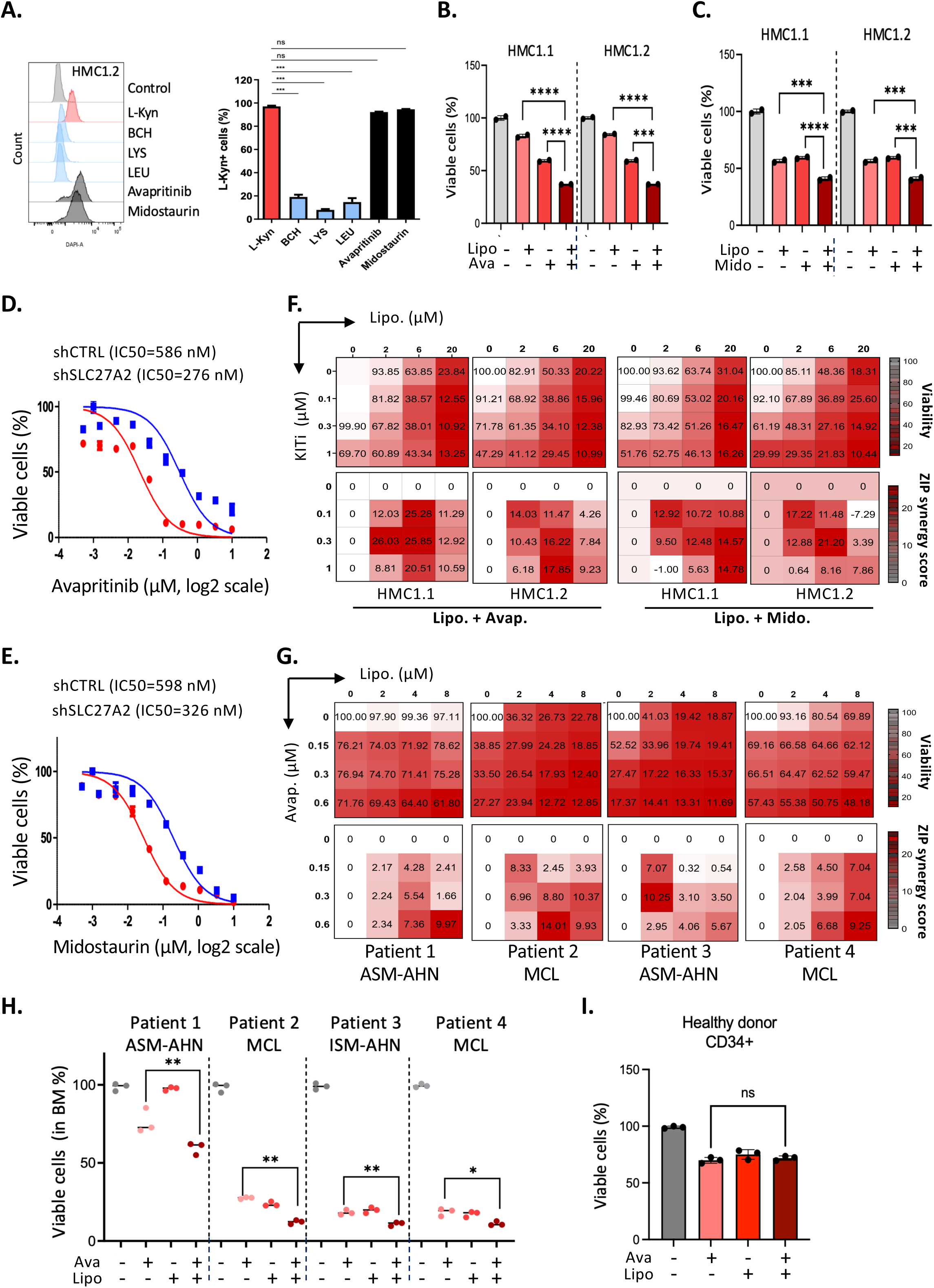
FATP2 inhibition sensitizes neoplastic mast cells to KIT-directed therapy. (A) L-kyn uptake by HMC1.2 cells in the presence or absence of the KIT inhibitors avapritinib or midostaurin inhibitors. Lysine (LYS) and leucine (LEU) were used as competitive amino acid controls for LAT-1-mediated L-kyn uptake, whereas BCH served as a positive control for System L transporter inhibition. (B-C) Effects of combined lipofermata and KIT inhibitor treatment (avapritinib or midostaurin) on neoplastic MC viability. (D-E) Genetic silencing of *SLC27A2* and its effect on neoplastic MC sensitivity to KIT inhibition, demonstrating a phenotype consistent with pharmacological FATP2 blockade. (F-G) Representative ZIP synergy plot illustrating the interaction between lipofermata and avapritinib across treatment matrices in neoplastic mast cell lines. (H) *Ex vivo* assessment of combination treatment activity in primary bone marrow cells from patient with AdvSM. (n=4) (I) Effect of combination treatment on CD34+ hematopoietic progenitor cells from healthy donors (n=3). Data are represented as mean ± SEM from at least three independent experiments, where applicable. *P < 0.05; **P < 0.01; ***P < 0.001; ****P < 0.0001; ns, not significant.

We next asked whether disrupting this parallel pathway could enhance the activity of KIT inhibitors. Sequential exposure to lipofermata (0.6 µM, 4 hours) followed by avapritinib or midostaurin (250 nM) resulted in a greater reduction in cell viability than either treatment alone in both HMC1.1 and HMC1.2 cells (Figures 6B-C). Genetic silencing of SLC27A2 reproduced this sensitizing effect (Figures 6D-E), supporting a specific contribution of FATP2 rather than an off-target pharmacological effect, and ZIP analysis in SynergyFinder 2.0 demonstrated synergy between lipofermata and each KIT inhibitor in both cell lines (Figures 6F-G).

Ex vivo, the enhanced activity of the combination was recapitulated in primary bone marrow cells from four patients with AdvSM (2 SM-AHN, 2 mast cell leukemia) (Figure 6H). At the concentration used in combination experiments (0.6 µM), lipofermata did not impair the viability of CD34+ progenitor cells from healthy donors (Figure 6I).

Kynurenine uptake and FATP2-dependent lipid acquisition therefore remain operational during KIT inhibition, and targeting this pathway enhances the activity of avapritinib and midostaurin.

## DISCUSSION

Although avapritinib and midostaurin have substantially improved outcomes in AdvSM, residual neoplastic MCs persist in many patients despite clinical responses ^8,9^ Recent single-cell analyses have confirmed the persistence of residual clonal populations under KIT inhibition.^34^ The non-genetic mechanisms enabling these cells to survive targeted therapy remain poorly understood. Here, we identify a kynurenine-AHR-FATP2 axis that promotes fatty acid uptake and membrane remodeling in neoplastic MCs, persists despite KIT inhibition, and whose disruption enhances KIT inhibitor activity in cell lines, primary patient cells, and *in vivo*, identifying lipid acquisition as a therapeutically actionable vulnerability in AdvSM.

Increased circulating kynurenine and reduced tryptophan levels have previously been reported in mastocytosis, primarily in relation to gastrointestinal and neuropsychiatric manifestations.^21,22^ Our findings link altered tryptophan metabolism to disease aggressiveness and neoplastic MC biology. Plasma L-kyn discriminated advanced from indolent disease in our cohort (AUC 0.94; n=25) and clustered with quinolinate, cholesterol, and GlcNAc, an oncometabolite that we previously associated with aggressive disease^23^, suggesting that kynurenine contributes to a circulating metabolic signature of AdvSM. Increased expression of IDO1, LAT1, AHR and ARNT in purified patient-derived MCs together with a coordinated tryptophan-AHR signature in a bone marrow compartment enriched for monocytes and MCs, indicates that this pathway is engaged within the neoplastic hematopoietic compartment. Kynurenine may therefore be not merely a systemic correlate of advanced disease but also an upstream signal influencing neoplastic MC metabolism. Although AHR is an established regulator of normal MC homeostasis,^52^ our findings suggest that it additionally functions as a metabolic sensor in neoplastic MCs.

The kynurenine-induced metabolic state, characterized by increased mitochondrial dependency and fatty acid oxidation, resembles the oxidative, lipid-dependent phenotypes associated with therapeutic resistance in acute myeloid leukemia (AML).^53–55^ AHR signaling has likewise been implicated in metabolic reprogramming in AML.^56^ Our findings extend this paradigm to a KIT-driven neoplasm. Kynurenine enters neoplastic MCs through System L amino acid transporters, including LAT1,^29^ and engages an AHR-dependent program associated with increased SLC27A2/FATP2 expression. FATP2 was the most highly expressed and most strongly upregulated fatty acid transporter in advanced disease. Although FATP2 has been characterized primarily in tumor-infiltrating neutrophils and myeloid-derived suppressor cells, where it supports immunosuppressive reprogramming,^45,57,58^ kynurenine-AHR signaling also contributes to these suppressive states.^59^ We extend this biology to neoplastic MCs, identifying FATP2 as a link between extracellular kynurenine sensing and lipid acquisition within the malignant cell itself.

The consequences of kynurenine-AHR signaling extend beyond bioenergetics: kynurenine exposure increased plasma membrane cholesterol, AHR knockdown reduced GM1-positive raft content, and combined cholesterol and sphingolipid depletion reduced KIT and STAT5 phosphorylation, consistent with the established requirement for lipid raft integrity in KIT-dependent survival and proliferation signaling.^47^ Increased lipid acquisition may therefore support malignant MC fitness through two complementary mechanisms: providing substrates for β-oxidation and maintaining the membrane organization required for oncogenic signaling. A key feature of this pathway is its persistence during KIT inhibition. Neither avapritinib nor midostaurin reduced kynurenine uptake, and both pharmacological and genetic FATP2 inhibition enhanced KIT inhibitor activity in KIT-mutant cell lines and in primary AdvSM bone marrow cells *ex vivo*. FATP2 inhibition therefore targets a metabolic vulnerability not directly addressed by current KIT-directed therapies, although its effects on lipid raft integrity suggest an additional, indirect impact on KIT-STAT5 signaling. Lipofermata also reduced disease burden in a KIT D816V xenograft model, was more active against neoplastic MC lines than against stromal and solid tumor cell lines, and did not compromise the viability of normal CD34+ progenitor cells viability at the concentrations used in combination. These findings support the feasibility of targeting FATP2 alongside KIT inhibition.

Plasma L-kyn is readily measurable and, together with GlcNAc, may contribute to a metabolic signature of aggressive disease. Because AHR nuclear translocation is regulated by O-GlcNAcylation^60^, the association between circulating kynurenine and GlcNAc raises the possibility that hexosamine flux modulates AHR activity in neoplastic MCs, suggesting an additional layer of crosstalk that warrants investigation.

The limitations of this study include the size of our metabolic cohort. Plasma L-kyn emerges as a promising biomarker candidate, whose discriminative value now awaits confirmation in larger, treatment-annotated cohorts in which systemic inflammation, renal function, and associated hematologic neoplasms can be modeled as covariates, as these factors may influence circulating L-kyn levels. The xenograft data establish *in vivo* proof of principle for FATP2 targeting independently of host immunity, leaving the immunological arm of this axis unexplored. Immunocompetent models and *in vivo* testing of the FATP2-KIT inhibitor combination represent the next steps toward clinical translation.

In summary, kynurenine-AHR signaling drives fatty acid acquisition and FATP2 dependency in neoplastic MCs. By operating alongside KIT and persisting despite its inhibition, this program sustains malignant cell fitness under targeted therapy. By linking extracellular metabolite sensing to lipid uptake, membrane organization, and KIT-associated signaling, our findings uncover a metabolic vulnerability in AdvSM and provide a rationale for combining metabolic targeting with KIT-directed therapies as a strategy to deepen responses and limit residual neoplastic MCs.

## Acknowledgments

The authors thank the patients and their families for participating in this study, and all the members of the French Reference Center for Mastocytosis (CEREMAST) for coordinating patient inclusion and sample collection. The authors are grateful to S. Granjeaud and G. Bidaut of the Integrative Bioinformatics (CIBI) platform, and the Cytometry, TrGET, and, Animal Facility platforms at the Centre de Recherche en Cancérologie de Marseille (CRCM) and the Cytometry and Cell Sorting platform at the Centre de Recherches en Cancérologie de Toulouse (CRCT) for technical support. The authors also thank Dr Yannick Hamon (CIML, Marseille) for advice on filipin experiments and Dr Jean-Emmanuel Sarry (CRCT, Toulouse) for insightful discussions.

This work was supported by the ARC Foundation (PJA #202223051 to F.B.), the Laurette Fugain Association (ALF 2022/06 to F.B.), and research funding from the Cancéropôle Provence-Alpes-Côtes d’Azur, Institut du Cancer, and Région Sud (to F.B). G.S. and C.L. were supported by doctoral fellowships from the French Ministry of Higher Education and Research; G.S. additionally received a fourth-year doctoral fellowship from the ARC Foundation. Team METAML (Sarry-Brenet) is a member of the OPALE Carnot Institute, The Organization for Partnerships in Leukemia.

## Authorship

G.S., C.L., J.A., S.L., A.S., L.M., R.C., and A.G. generated data and performed statistical analysis; A.S. and L.M. performed Cholera toxin B staining and image analysis; R.C. and A.G. performed NSG mouse xenotransplantation experiments; L.MC., R.J.A, M.A. O.H., J.R., Y.C., A.O., and P.D. provided cell lines, SCENITH antibodies, patient samples, and clinical data; PK and PKK performed the bioinformatic analyses; G.S. and C.L. drafted the initial version of the figures; F.B. conceived and supervised the study and wrote the manuscript, which was critically reviewed by all authors. All authors read and approved the final manuscript.

## Data sharing statement

RNA sequencing data generated in this study are deposited in the Gene Expression Omnibus under accession GSE353477. Other data supporting the findings are available in the manuscript and its supplemental materials, or from corresponding author on request. Publicly available datasets are cited in the manuscript.

