## Supplemental Figures for "Kynurenine-AHR signaling sustains a targetable FATP2 dependency in advanced systemic mastocytosis"

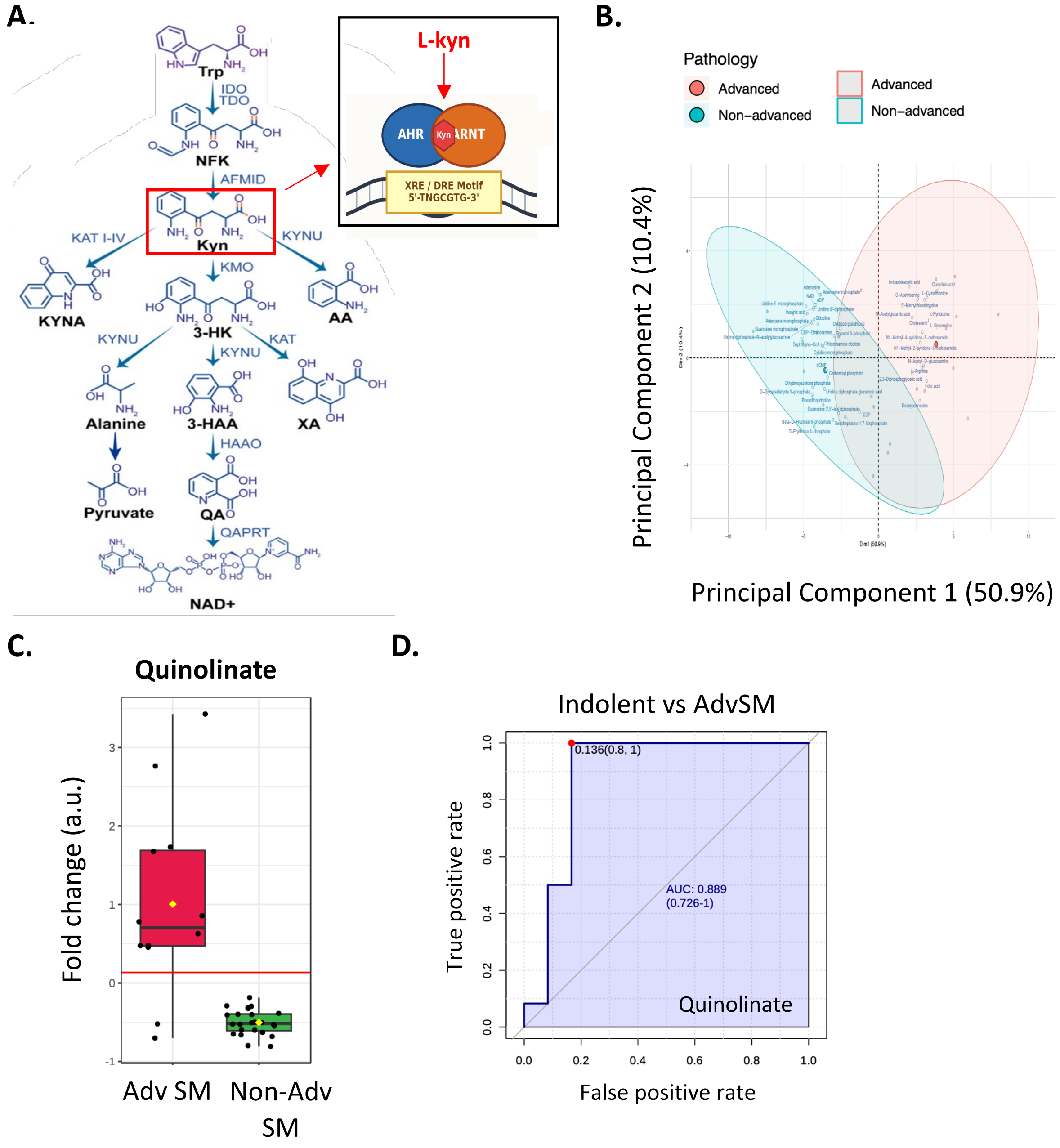


**Figure S1: Additional metabolomic analyses support enrichment of the kynurenine pathway in advanced systemic mastocytosis**

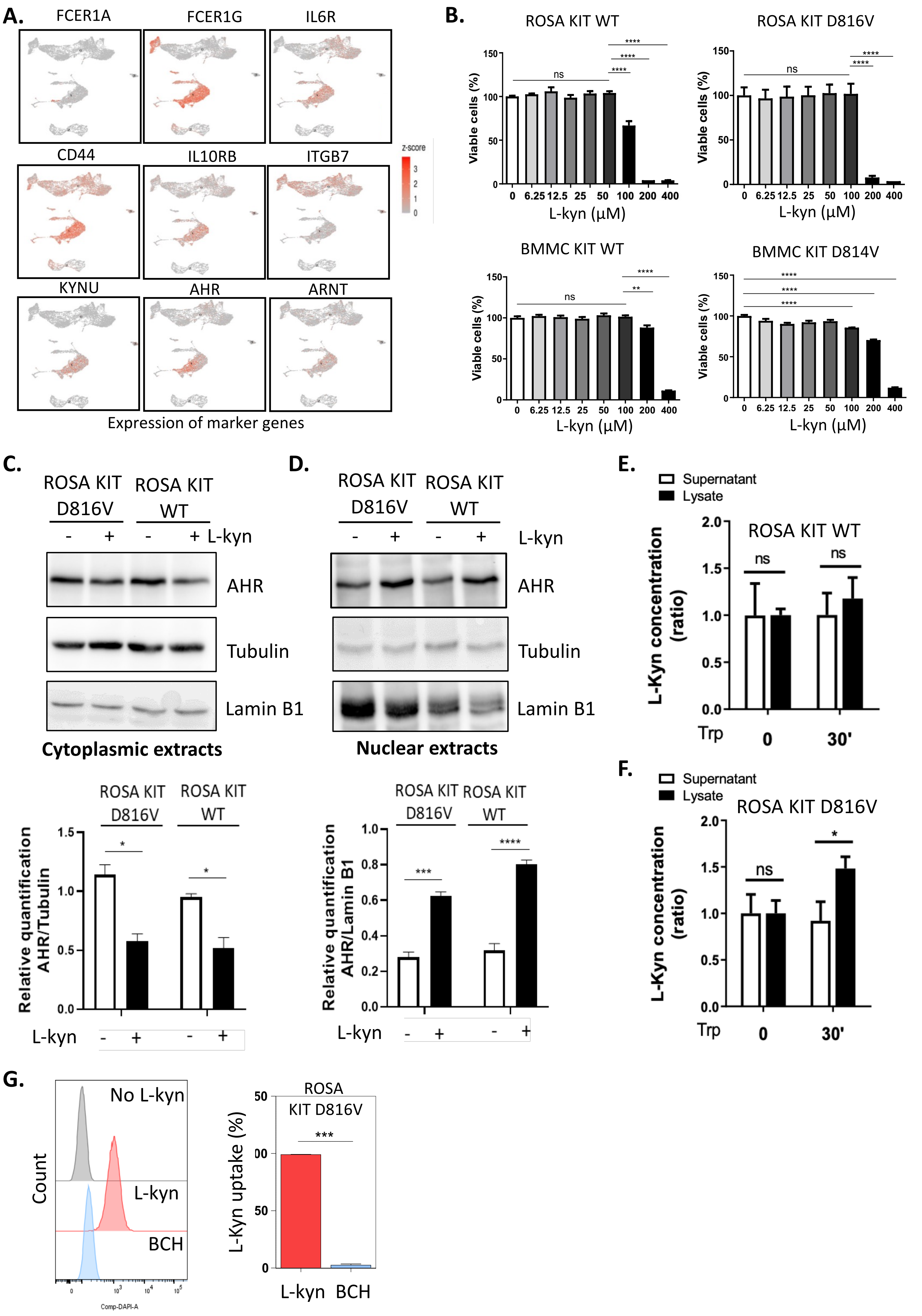


**Figure S2: L-kynurenine uptake induces AHR nuclear translocation in mast cell models.**

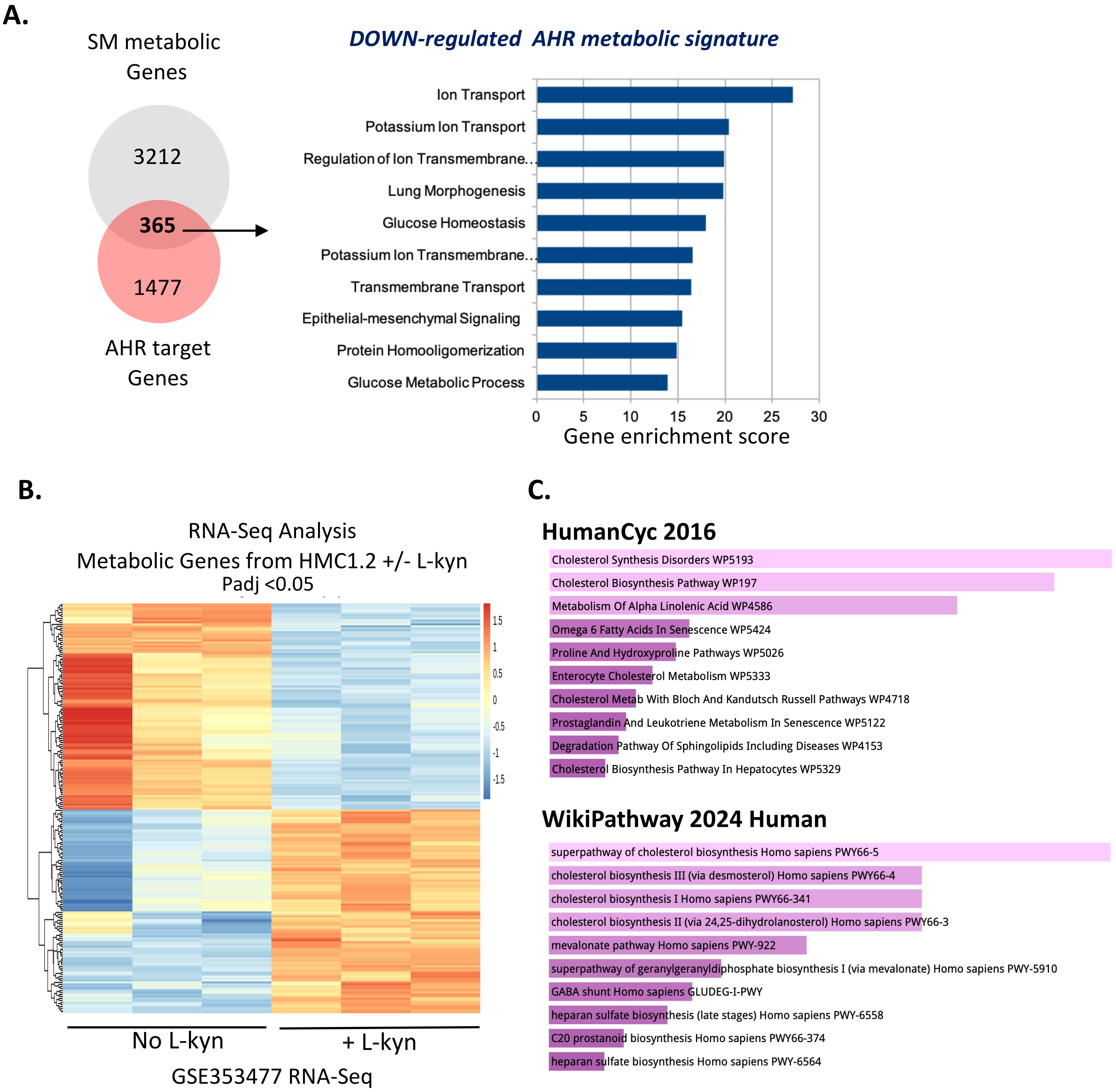


**Figure S3: Transcriptomic analyses define an AHR-linked lipid metabolic program in neoplastic mast cells.**

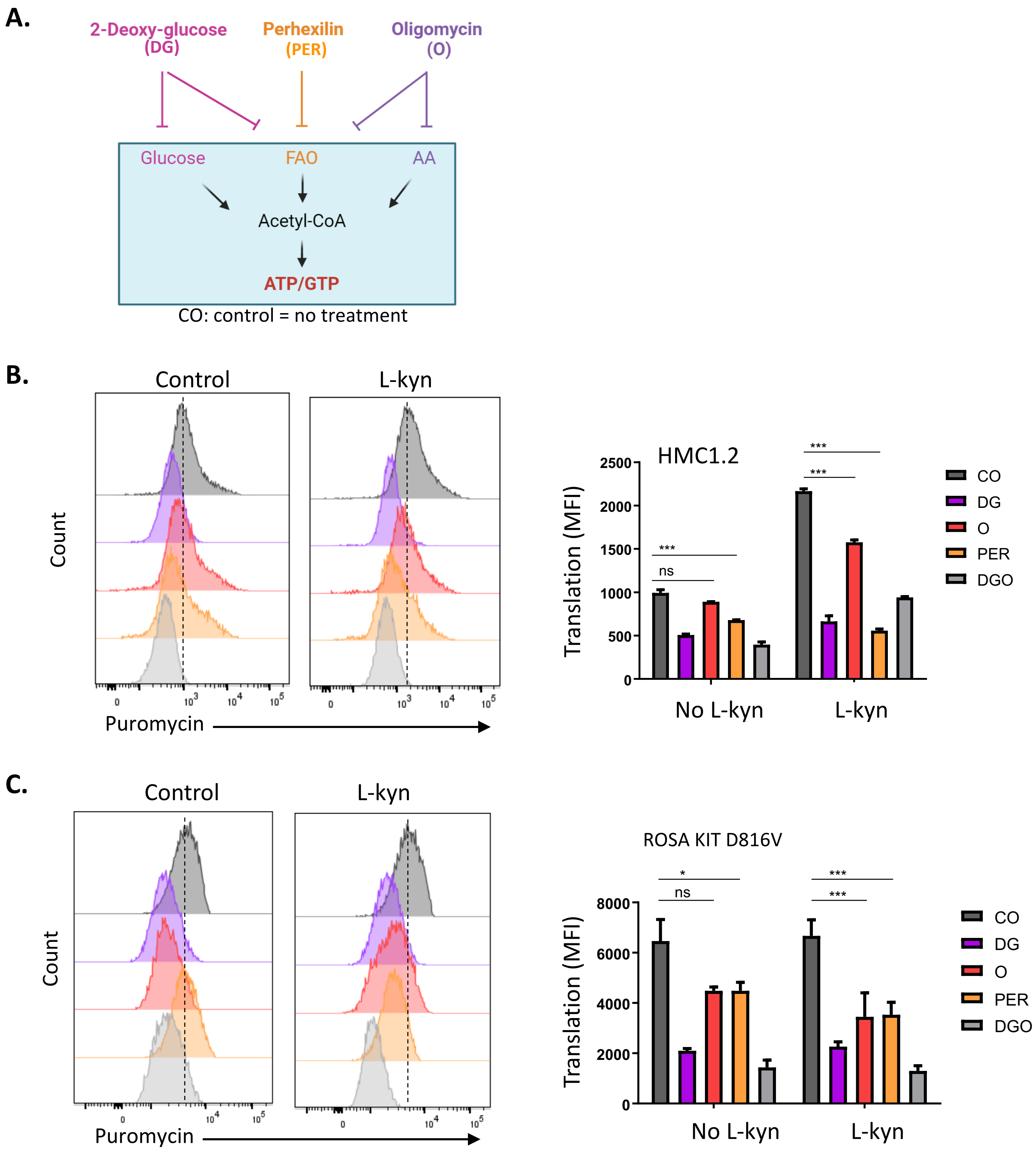


**Figure S4: L-kynurenine exposure alters translation-related metabolic profiles in neoplastic mast cells: SCENITH raw data.**

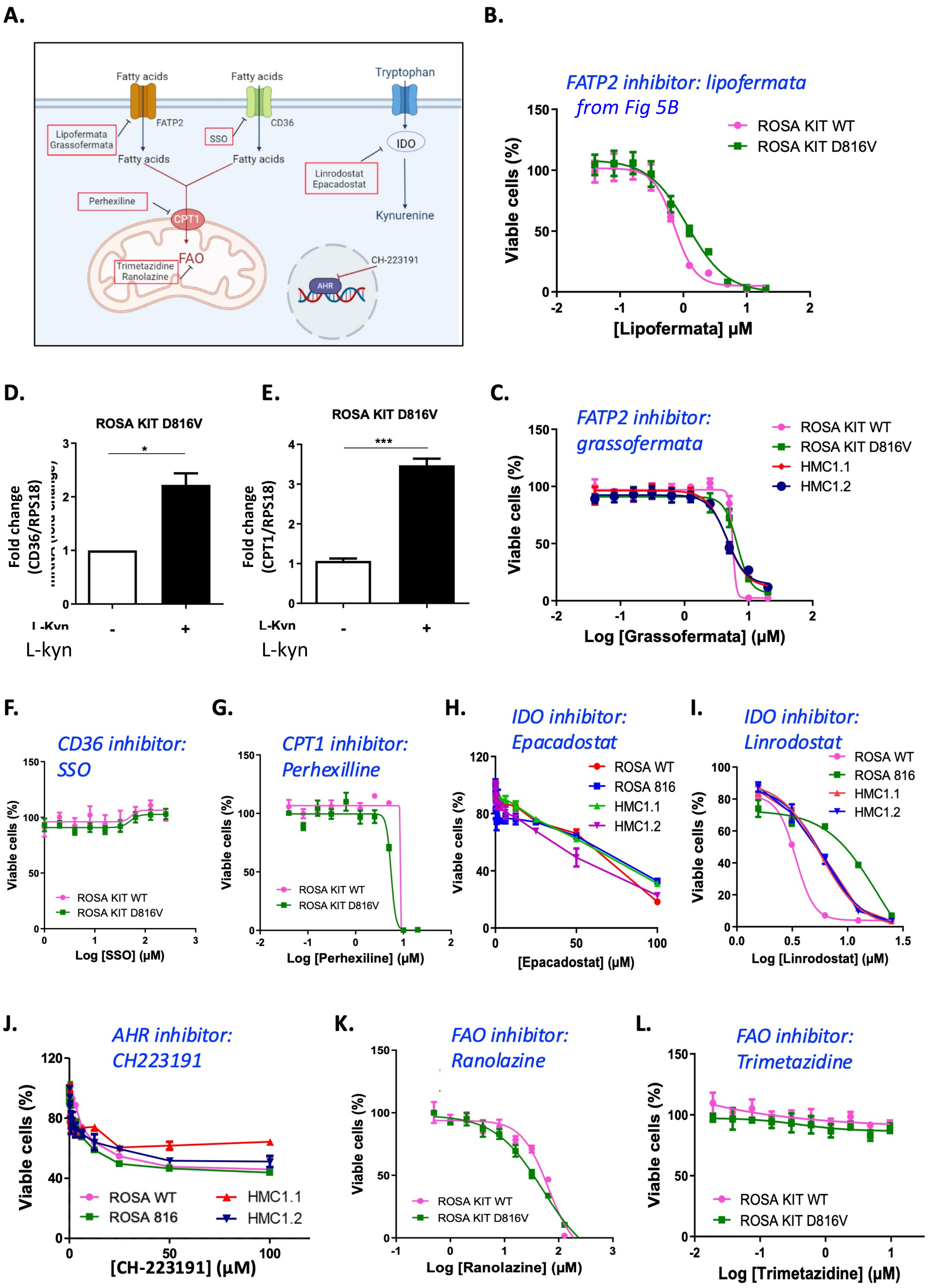


**Figure S5: Pharmacological inhibition of fatty acid oxidation at multiple levels impairs neoplastic mast cell viability.**
